# CellMAPtracer: A user-friendly tracking tool for long-term migratory and proliferating cells

**DOI:** 10.1101/2020.06.15.152462

**Authors:** Salim Ghannoum, Kamil Antos, Waldir Leoncio Netto, Alvaro Köhn-Luque, Hesso Farhan

## Abstract

**Background:** Cell migration is a fundamental cell biological process of key importance in health and disease. Advances in imaging techniques have paved the way to monitor cells motility. An ever-growing collection of computational tools to track cells has improved our ability to analyze moving cells. However, few if any tools let the user supervise and correct cell tracks that are automatically detected. Thus, we developed CellMAPtracer, a tool to track cells in a semi-automated supervised fashion, thereby improving the accuracy and facilitating the long term tracking of migratory and dividing cells. CellMAPtracer is available with a user-friendly graphical user interface and does not require any coding or programming skills.

**Results:** We used CellMAPtracer to track fluorescently-labelled BT549 breast cancer cells. It allowed us to track dividing cells and determine the fate of the daughter cells with respect to migration speed or directionality and cell cycle length. Of note, we were able to track multi-daughter divisions, wherein a cell divides and gives rise to more than two cells. We also identified a not previously described speed change in the terminal phase of the cell cycle.

**Conclusion:** CellMAPtracer is a software tool for tracking cell migration and proliferation through a user-friendly interface that has a great potential to facilitate new discoveries in cell biology.

## Background

Cell migration is a fundamental biological process that plays important roles during tissue morphogenesis, immunology, wound healing, or tumor progression (Friedl and Gilmour, 2009). Thus, investigating cell motility is important for many research areas such as developmental biology, physiology, neuroscience, or cancer biology. The visualisation of fluorescently-labeled cells using time-lapse video microscopy experiments plays an important role in understanding how cells move and what governs their migratory movements. To draw robust conclusions, cell movements need to be tracked with a high level of accuracy and a minimum level of bias. Thus, the development of tools for automated or semi-automated cell tracking is key to obtain quantitative data from biologic experiments. However, identifying and tracing fluorescently-labeled cells that migrate, proliferate, interact or die from time-lapse microscopy is a laborious and error-prone process.

Historically, cell tracking has been performed by manual selection on a reference point within a cell for each time-lapse frame (Masuzzo et al., 2016). While workable in occasions, see for instance (Sugihara et al., 2015), this approach is often prohibitively time consuming and prone to user bias due to the difficulty in manually defining cell positions and the noticeable inconsistency between different users (Cordelières et al., 2013). Manual tracking is also error-prone to either over-representing particular angles between displacements or repeating previous x-y coordinates (Gorelik & Gautreau, 2014). On the other hand, automated tracking algorithms can provide objective migration tracks due to the elimination of errors associated with the human factor. However, automated tracking can itself generate artifacts due to the high possibility of undergoing a flip-flop switch between neighbour cells within the track of a particular cell. Thus, the tracking efficiency decreases dramatically with high density and fast movements. This may obscure biologically-relevant differences between experimental settings or generate spurious results. Today, a large number of cell tracking systems exist (Sacan et al., 2008; Shen et al., 2010; Van Valen et al., 2016; Cooper et al., 2017; DuChez, 2017; Tsai et al., 2019). Unfortunately, the majority of automated tracking tools lack track supervision and editing functionality features, such as target selection, trajectory inspection and error correction (Meijering et al., 2012). The absence of these features requires adjusting numerous parameters to optimize performance to the point where tracks are acceptable. Furthermore, the difficulty level in automated tracking depends on the quality of the recorded video sequences. Often, time spent on adjusting parameters and recalculating cell tracks is comparable with full manual tracking. Importantly, tracking moving proliferating cells in time-lapse video sequences remains challenging due to the change in the morphology and the size of the labeled cells/nuclei. Many tracking systems are not sensitive to cell division but they can only track cells within one cell-cycle period. Some systems can keep tracking cells for more than one generation by selecting one of the two daughter cells. In both cases, all tracked cells are originally independent with indistinguishable cell division periods throughout the track. Therefore, studying the migratory changes during cell division is not applicable.

To solve these problems, we introduce CellMAPtracer, an open, free and user-friendly graphic interface for tracking fluorescently-labeled cells. CellMAPtracer allows studying migratory and proliferating cells in a supervised and semi-automatic fashion. CellMAPtracer is applicable for a variety of 2D cell migration assays, such as random migration and directed migration. It is capable of combining automated tracking with manual curation. It provides basic motility analysis and categorized trajectory data for deeper trajectory investigation. CellMAPtracer allows users to trace and follow individual cells throughout the course of the live-imaging. This can enable the user to visualize the tracks of the descended cells and their ancestor in an interactive multi-generation plot. The obtained trajectory data can be used to precisely estimate the doubling time of the tracked cell population as well as answering questions that are difficult to date. For instance, do two daughter cells migrate and divide synchronously? Does the cell directionality profile change before and during cell division? CellMAPtracer is the only user-friendly multi platform tracking software that allows precise long-term tracking of proliferating cells by simple inspection and correction (see Supplementary table 1).

## CellMAPtracer availability and Installation

CellMAPtracer is built using MATLAB (v 9.8) and can be freely obtained as: (A) a standalone executable program for Microsoft Windows, macOS or GNU/Linux; (B) a MATLAB App/Toolbox; and (C) the source MATLAB code.

### (A) CellMAPtracer standalone executable program

Navigate to https://github.com/ocbe-uio/CellMAPtracer/releases/tag/v1.0. Three assets of CellMAPtracer for Windows, Linux and macOS versions can be found and downloaded. After downloading the version compatible with your Operating System, users should uncompress the file and follow the instructions in the corresponding “readme.txt”.

### (B) CellMAPtracer MATLAB App

To be able to run CellMAPtracer App within the MATLAB environment, users should follow three simple steps: 1) Download the “App” folder from the CellMAPtracer repository: https://github.com/ocbe-uio/CellMAPtracer.2) In MATLAB, go to APPS tab and click “Install App” and find “CellMAPtracer.mlappinstall” then install it. 3) Open CellMAPtracer App from Application list in MATLAB.

### C) CellMAPtracer from the source MATLAB code

To be able to run CellMAPtracer code, users should clone the CellMAPtracer repository from https://github.com/ocbe-uio/CellMAPtracer and then run “CellMAPtracer_Main.m” after opening a project in MATLAB.

## CellMAPtracer description & workflow

CellMAPtracer is a desktop application with a graphical user interface (GUI) capable of loading multi-TIFF stacks (8 and 16 bits) of spatio-temporal live cell images as input (Fig. 1a). The output is an interactive multi-generation trajectory plot (Fig. 1b) and 5 categories of trajectory data. The 5 categories include: all cells, dividing cells, non-dividing cells, daughter cells and dividing daughter cells. Each of these contains two excel sheets. The first sheet contains the measurements of cell migration parameters such as the total distance, displacement, directionality and speed. The second sheet contains the x-y coordinates of tracked cells in the corresponding category (Supplementary Fig. 1). The purpose of the categorization is to enable users to easily plot the cell migration statistics without the need of advanced programming skills. Such plots can help highlight the migratory phenotype of cells in each category and draw conclusions about the doubling time, the heterogeneity of daughter cells, speed-directionality dynamics prior and through cell division.

**Fig. 1:**
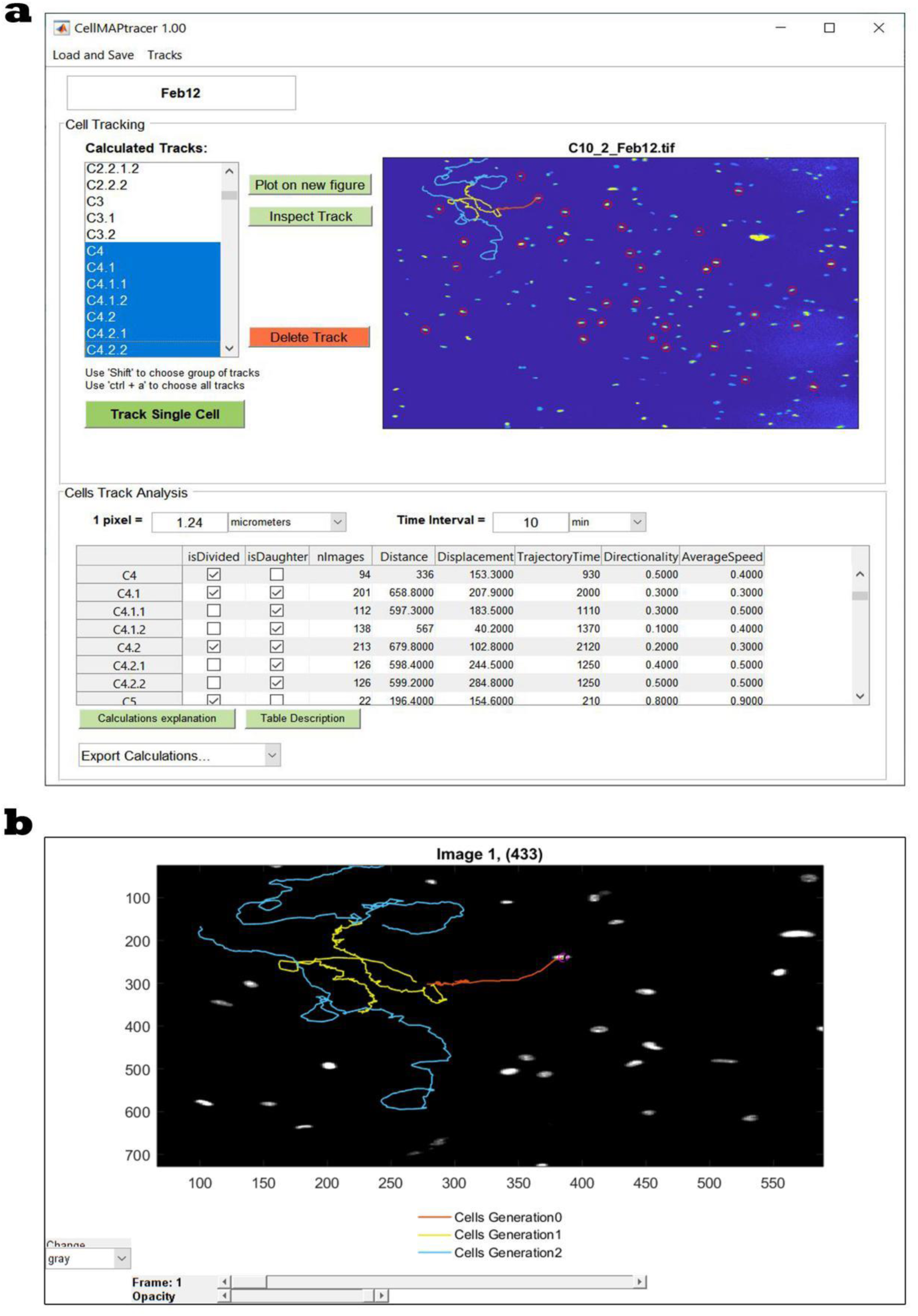
CellMAPtracer graphical user interface. (a) The main user interface window used to track a single cell and all its descendant cells. (b) A representative interactive multi-generation trajectory plot of a cell (orange) and its descendant daughters (yellow) and granddaughters (blue).

CellMAPtracer is based on two detection approaches for nuclei segmentation. The first approach uses a paradigm called tracking-by-detection. It relies on a fluorescence detector to initialize, adjust, reinitialize, supervise, and terminate a tracker (Breitenstein et al., 2011). CellMAPtracer analyses the frame-to-frame position of a target cell. For each frame, contrast-limited adaptive histogram equalization is used to separate nuclei from the background by converting greyscale images into binary images. All above-threshold contiguous regions are considered nuclear objects taking into account the spatial characteristics of segmentation. The second approach of cell segmentation is the watershed transformation. This finds ‘watershed ridge lines’ in an image and treats an image as a surface where light pixels of the nuclei represent high elevations and dark pixels of the background represent low elevations (Meyer, 1994). This fast and intuitive method allows to separate close nuclei from each other regardless the similarity degree in the signal intensity (Kornilov & Safonov, 2018). As a result, highly accurate, instance segmentation is generated. After nuclei detection, the position of each nucleus is determined by finding its centre of mass, which is calculated based on the basis in all of all the pixels in the particle having the same intensity. Positions of all nuclei within the field of view are compared with the position of the tracked cell on the last frame. To find the new cell position, the algorithm calculates the distances between the last position of the target cell nucleus and current position of each other nucleus. The position with minimum distance is set as the new position of the tracked cell. (Supplementary Fig. 2). To save computational time, the local features are computed only within a fraction of image defined by the interactive slider in the CellMAPtracer tracking window.

**Fig. 2:**
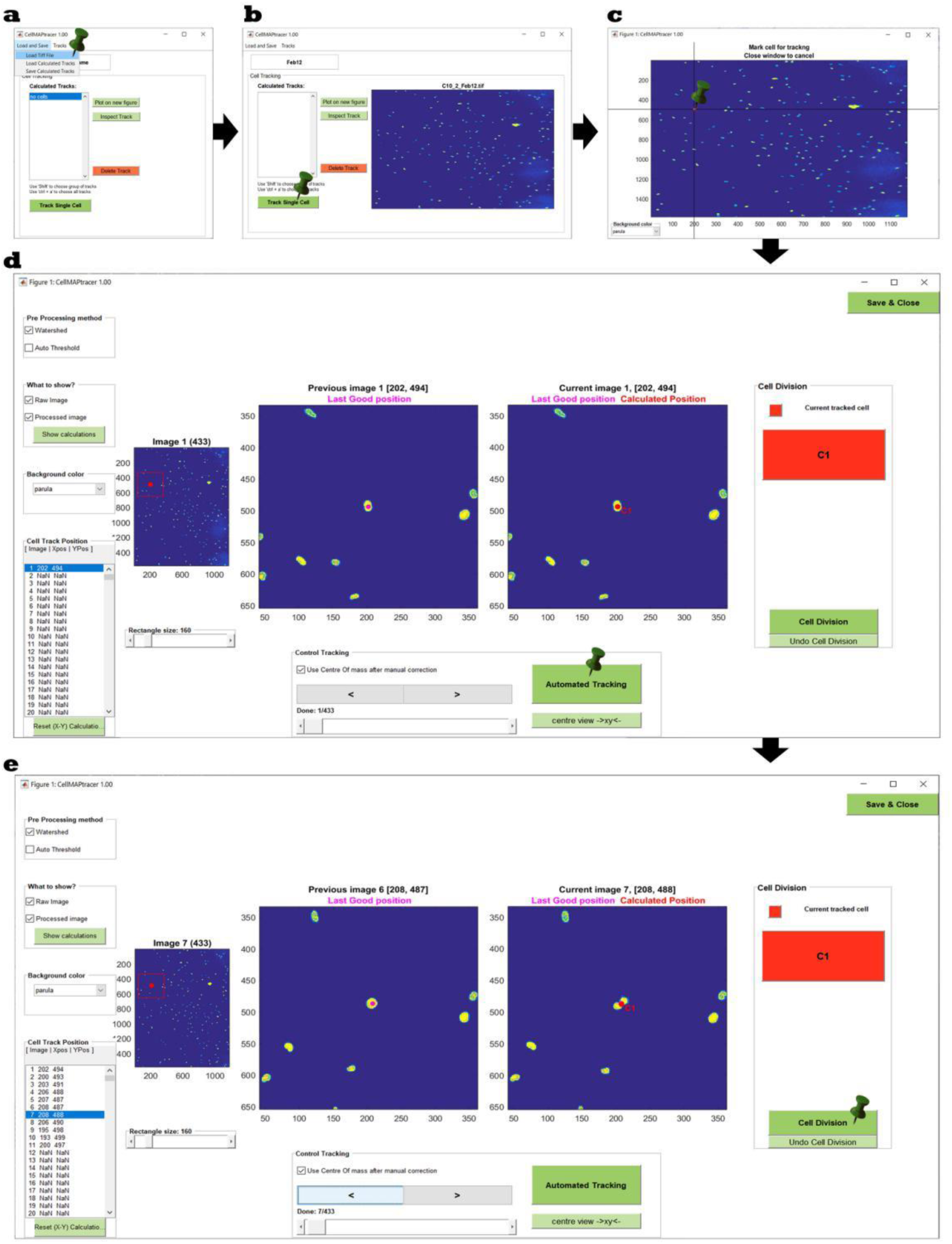
CellMAPtracer workflow. (a) CellMAPtracer front window, where users can load either a TIFF image stack or calculated tracks. (b) Starting the tracking process by clicking on “Track Single Cell” button. (c) Selecting the cell to be tracked. (d) Tracking window, where users can run the automated tracking and monitor it step-wisely. (e) Marking a cell division by clicking on the “Cell Division” button.

We designed CellMAPtracer to allow cell tracking to be done automatically and monitored step-wisely (Fig. 2). Tracking errors can distort the cell trajectory results. For that reason, we designed CellMAPtracer interface to include multi-features for importing, highlighting cell division, inspecting and correcting existing tracked cells/nuclei to reach near 100% tracking accuracy. In case the algorithm mistakenly tracked the target cell or switch to another cell, the user can manually correct the tracking by pausing the automated tracking and clicking on the correct position on the current frame. The corrected position will be corrected further if the “Use center of mass after manual correction” button is activated. Otherwise the exact position selected by the user will be recorded. In case the intensity of the target cell/nucleus is very low, the “Use center of mass after manual correction” button should be deactivated.

During the course of tracking, the migratory cell might undergo a cell-division. CellMAPtracer allows marking cell divisions and lineage tracing of all descendant cells independently. From intensity and morphology perspectives, cell division is a very dynamic process. The naked eye can in most cases recognize the occurrence of cell division. However, the automated cell division detection, which is mainly benefited from the representational power of deep learning models, requires enormous computational power and training copious amounts of data (Ciaparrone et al., 2020). To handle such difficulties, when the user notices a division event a single-click user intervention is needed. This simple intervention initiates tracks for the new daughter cells. There is no limitation of number of marked divisions. Therefore, CellMAPtracer can optimally handle long-term live imaging experiments.

## Results

As a proof of concept, we use CellMAPtracer to track and analyze human breast cancer migrating randomly on a 2D space. We used the triple-negative breast cancer cell line of BT549, stably expressing nuclear green fluorescent proteins (GFP). The methodology of the cell culture, the generation of GFP-BT549 stably-labeled cells, the dense-random migration assay and live imaging are explained in the supplementary materials (supplementary file S1).

From the first frame of three multi-TIFF, 8 bits image stacks (Ghannoum & Antos, 2020), a total of 103 cells were manually selected to be tracked. Other cells were also tracked but they were manually excluded due to fluorescence intensity issues or early disappearance from the scanning field. These 103 cells and their descendants were followed during 72 hours of live imaging. At the end of the tracking course, a total of 648 cells were tracked. From all tracked cells, 42% underwent a cell division. A 27% of all tracked cells were dividing daughter cells, which are generated from parent cells and undergone a second round of division themselves. The trajectory time of those dividing daughter cells gives a precise estimation of the doubling time. In our case, the doubling time of BT549 cells (n=175) averaged 31.1 ± 8.5 hours with a median of 30.2 hours and a mode of 24.7 hours (fig. 3a-3b). Correlation analysis showed a negligible correlation between the doubling time and both the speed and directionality of movement (fig. 3c). On the other hand, the doubling time was moderately correlated (r = 0.6) with the total distance (fig. 4d).

**Fig. 3:**
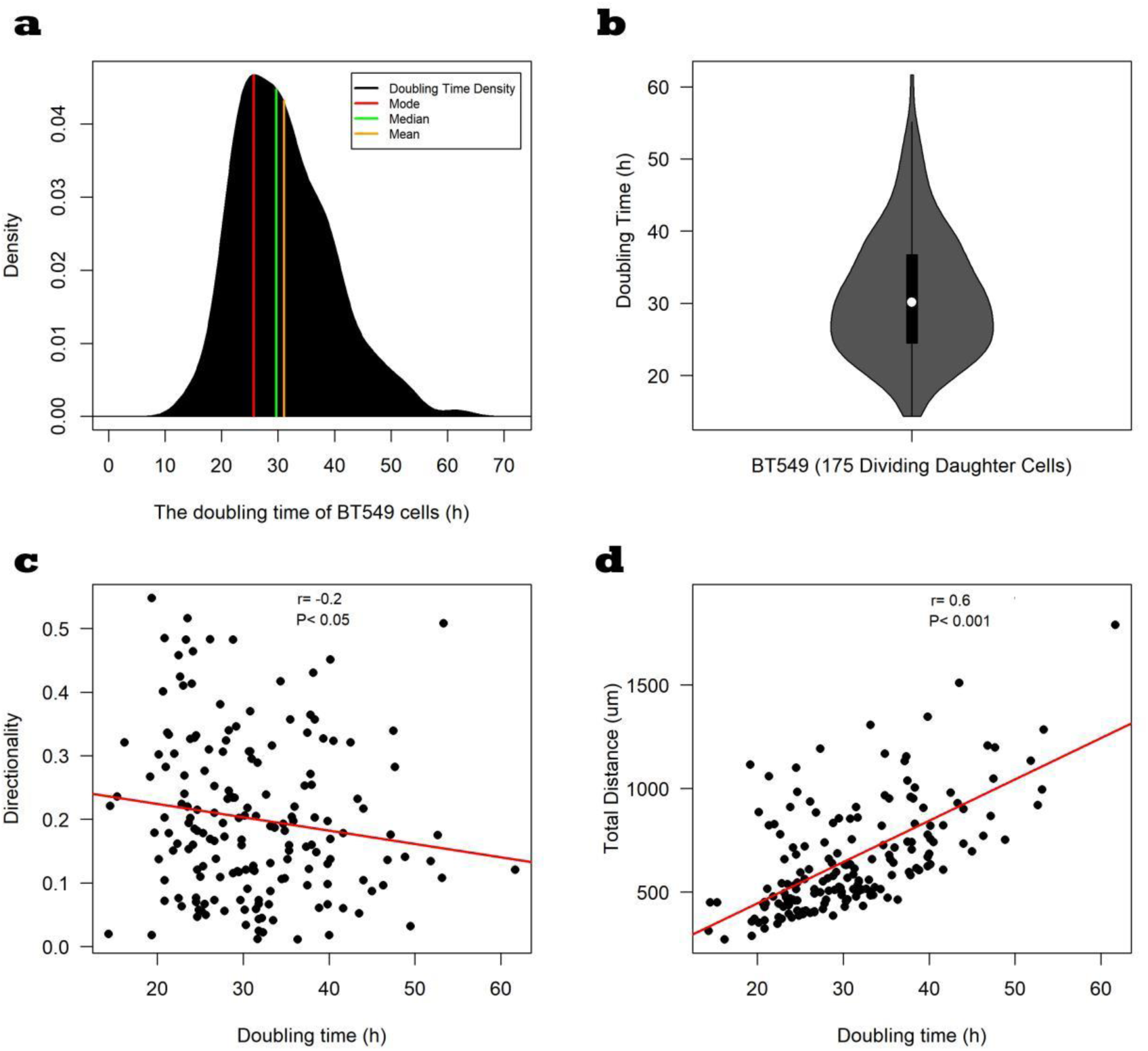
Descriptive and correlation statistics of the doubling time of BT549 cells (n=175). (a) Density plot of the doubling time of BT549 cells. (b) Violin plot showing the median and the interquartile range of the doubling time of BT549 cells. (c) Very weak negative correlation (Spearman’s rank correlation coefficient= -0.2) between the doubling time and directionality. (d) Moderate correlation (Spearman’s rank correlation coefficient= 0.6) between the doubling time and the total distance.

**Fig. 4:**
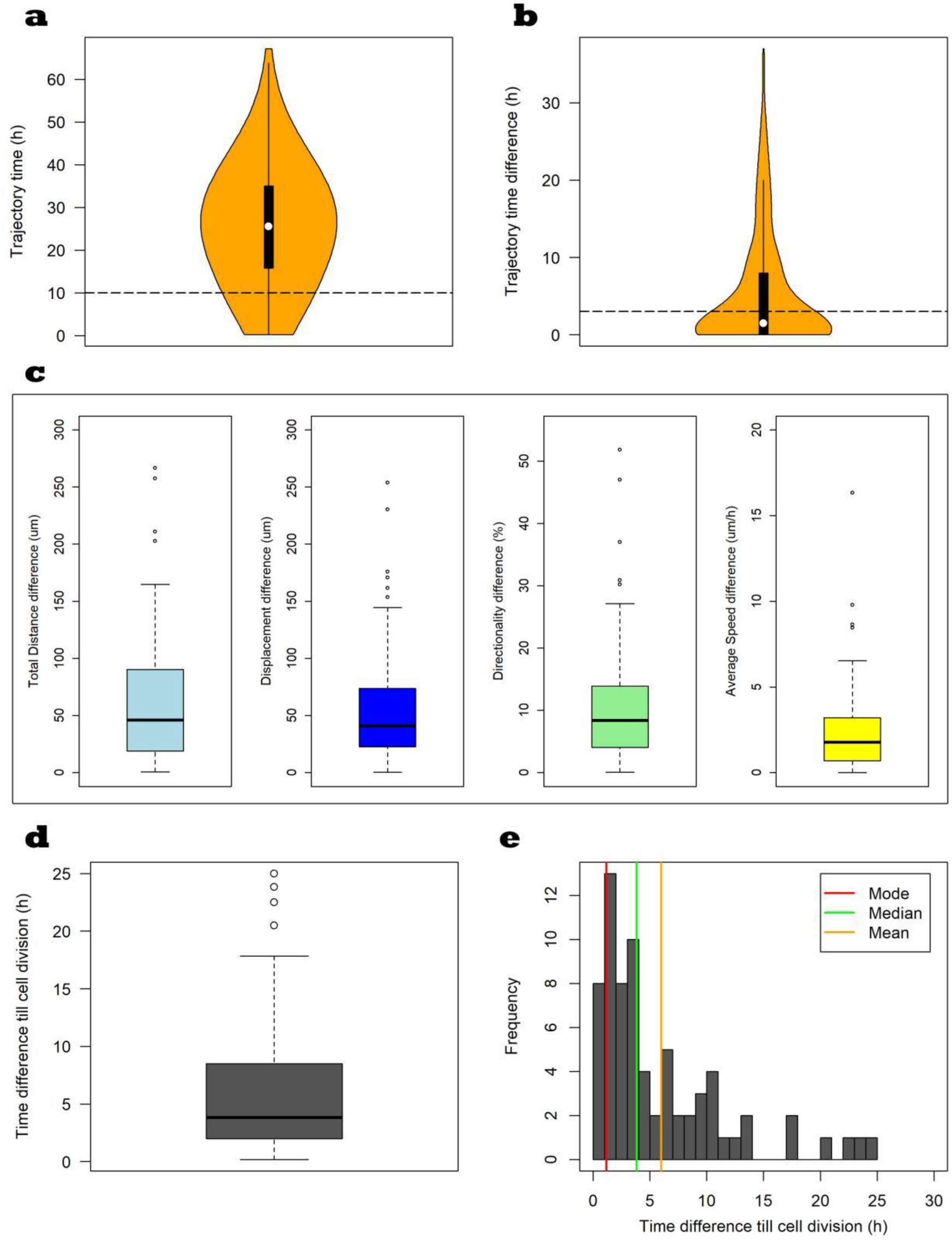
Heterogeneity between BT549 daughter cells. (a) Violin plot showing the median and the interquartile range of the trajectory time of daughter cells (n=540), the dashed line refers to the minimum required trajectory time of 10 hours. (b) Violin plot showing the distribution of the trajectory time difference between daughter cells (n=540), the dashed line refers to the maximum allowed trajectory time difference of three hours. (c) Boxplots showing the heterogeneity between of BT549 daughter cells (n=120 pairs) based on 4 migration measures of total distance, displacement, directionality and average speed. (d-e) The distribution of the time difference between daughter cells (n=71 pairs) till cell division.

Next, we show how CellMAPtracer enables users to gain insights about the heterogeneity between daughter cells. For a meaningful comparison, the considered trajectory time should be long enough and comparable between the two daughter cells. Here, and based on the distribution of the trajectory time of daughter cells (Fig. 4a-4b), we selected the cells that have a minimum trajectory time of 10 hours with no trajectory time difference larger than 3 hours between the two daughter cells. A total of 120 pairs of daughter cells were used as input for the heterogeneity analysis. Interestingly, the two daughter cells show relatively different trajectory measure values (Fig. 4c). The median difference in the total distance is 46.00μm. The median difference in the displacement is 40.90μm. The median difference in the directionality is less that 10%. The median difference in the average speed is 1.77μm /h. We also show how to analyse the synchronization degree in undergoing cell division. For that, only daughter cells with both cells undergoing cell division were included in the analysis (n=70 pairs). This can be easily done by selecting the cells in the “Daughter cells excel file” based on the “isDivided” column. It is noticeable that the majority of the two daughter cells do not divide at exactly the same time, but with some difference in the range of 5 hours (Fig. 4d-4e). Only 10% of the daughter cells divided with precise synchronization. The mode time difference is 1.2 hours. The mean time difference is 5.7 hours with a standard deviation of 5.5 hours. The time difference until cell division between daughter cells is weakly correlated with both the directionality difference (r=0.25) and the average speed difference (r=0.23) between the daughter cells (Fig. 5).

**Fig. 5:**
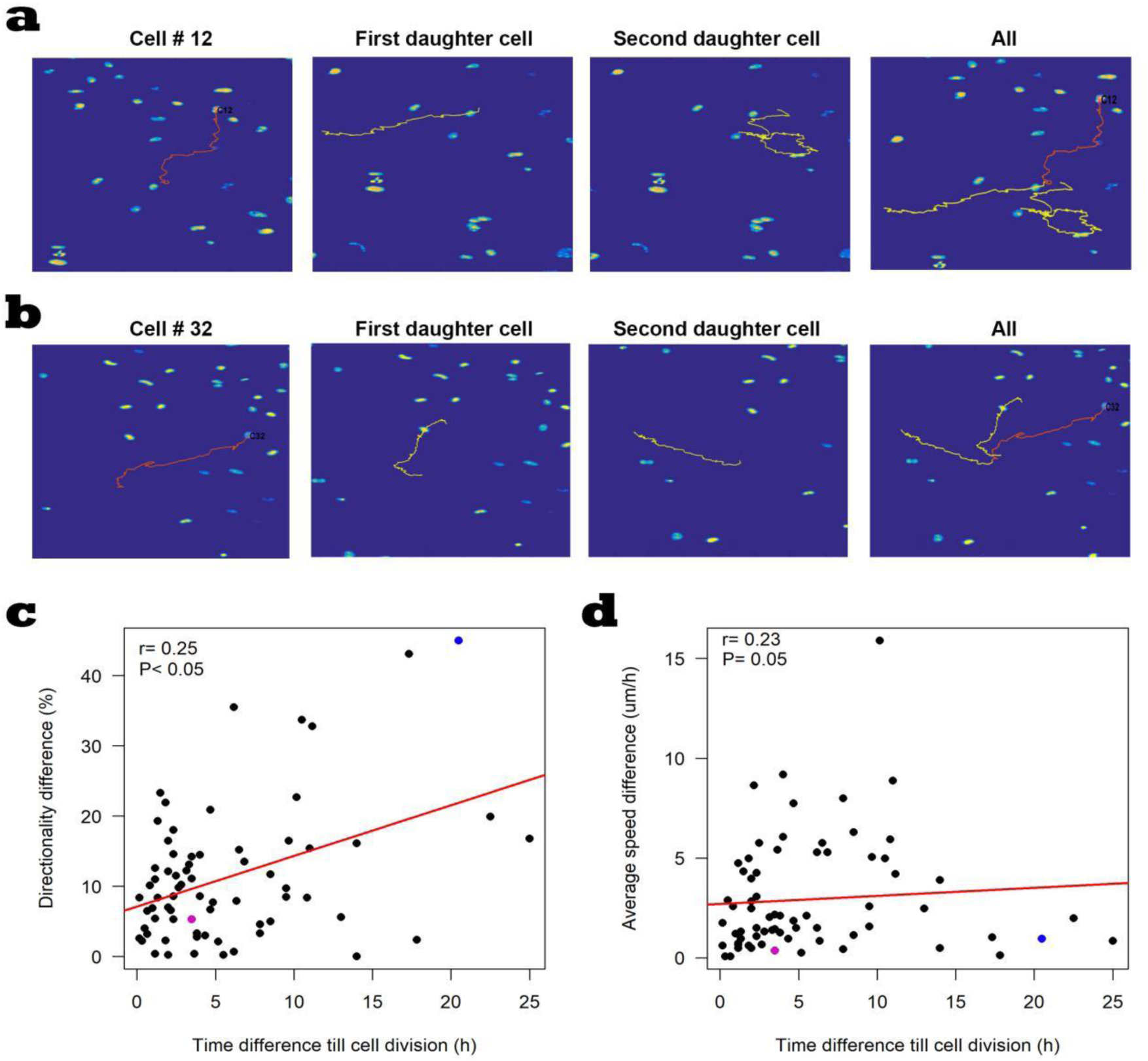
Representative examples of the heterogeneity between BT549 daughter cells. (a) An example of a cell (cell number 12 in red) with heterogeneous daughter cells (yellow). (b) An example of a cell (cell number 32 in red) with relatively homogeneous daughter cells (yellow). (c-d) Weak correlation between the time difference till cell division with both the directionality difference daughter cells (Spearman’s rank correlation coefficient= 0.25) and the average speed difference (Spearman’s rank correlation coefficient=0.23) between the daughter cells. The blue dot represents cell # 12 whereas the magenta dot represents cell # 32.

In order to study the speed-directionality dynamics prior and through cell division, cells with a minimum trajectory time of 24 hours and a maximum of 48 hours were selected (n= 152). The trajectory paths of the selected cells were analyzed for 24 hours prior generating daughter cells as a consequence of cell division. Moreover, the trajectory path was divided into 2 main phases, a preparatory phase of 20 hours and a G2-M phase of 4 hours (Cooper, 2000; Alberts et al., 2002). During the preparatory phase, the cell undergoes normal growth processes while also preparing for cell division. It consists of three stages; G1, S and the first half of G2. To be able to draw conclusions, we divided the preparatory phase into 5 equal periods (p1, p2, p3, p4 and p5), each of which has a trajectory time of 4 hours, which is similar to the trajectory time of the G2-M phase. The G2-M phase refers to a particular period of cell division, which includes the second half of the G2 phase and mitosis. From a cell migration point of view, the G2-M phase is highly important due to the disassembly of the Golgi apparatus (Rabouille & Kondylis, 2007). Intactness of the Golgi is known to play an important role in regulating cell migration (Bisel et al., 2008; Wei & Seemann, 2017; Millarte & Farhan, 2012). For the sake of simplicity, we will refer to the p5 period as the G2-M phase. The speed-directionality dynamics were analyzed for all the 152 cells. This was done by computing the migration measures of average distance per step, directionality and average speed in the preparatory and the G2-M phases as well as in each of the 5 periods of the preparatory phase. During the G2-M phase, the average distance per step and average speed of cells are significantly lower than corresponding measures during the first three periods of the preparatory phase (Fig. 6a and 6c) as well as lower than the average values of all the periods of the preparatory phase (Fig. 6b and 6d). Moreover, the directionality of the cells during the G2-M phase is significantly lower than their directionality during the preparatory phase (Fig. 6e-6f).

**Fig. 6:**
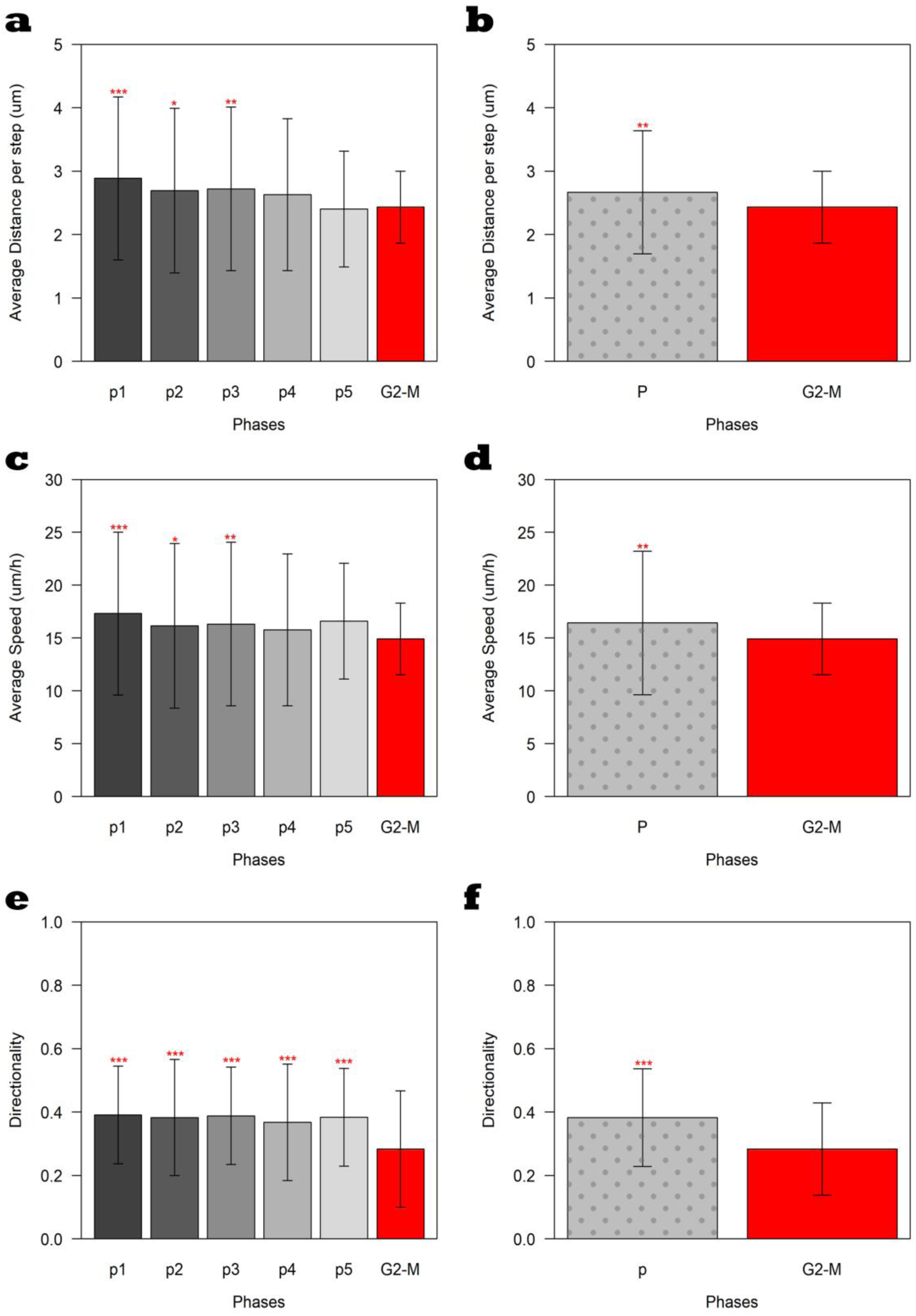
Migration measures in the preparatory and the G2-M phases (n=152). Gray bars represent the corresponding migration measure in the preparatory phase. The doted gray bars represent the average value of the corresponding migration measure in all the periods of the preparatory phase. Red bars represent the corresponding migration measure in the G2-M phase. (a-b) Bar plot of the average distance per step. (c-d) Bar plots of the average speed. (e-f). Bar plots of the directionality. (*P≤ 0.05; ** P≤ 0.01; ***P≤ 0.001)

Finally, we demonstrate that CellMAPtracer helps making discoveries in cell biology. During the tracking process we observed a phenomenon that, to the best of our knowledge, has not been previously described and that we propose to call “terminal speed jump”. The trajectory analysis of the tracked cells shows that 60.5% of the BT549 cells exhibit one or more instantaneous dramatic change in the speed within the last hour just prior generating daughter cells (Fig. 7). A cell is marked as undergoing a terminal speed jump if it shows one or more instantaneous speeds, during the last hour of G2_M phase, with at least double of the average instantaneous speeds during the whole G2_M phase. Moreover, that instantaneous speed should be at least three fold the neighbour instantaneous speed during the last hour of G2_M phase (Supplementary Fig. 3).

**Fig. 7:**
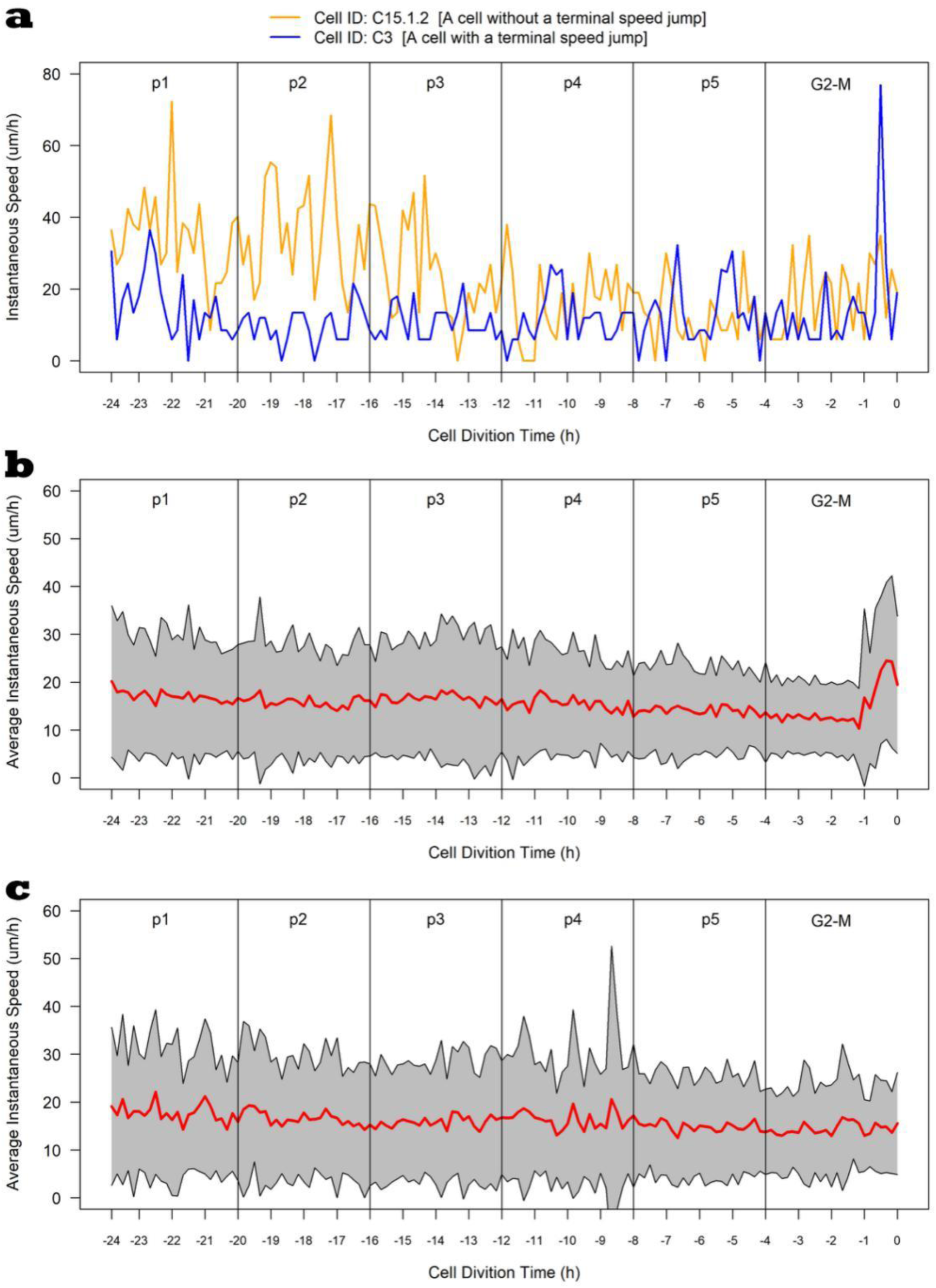
Speed Profile during the preparatory and the G2-M phases. (a) Examples of cells with a terminal speed jump [C3] and without [C15.1.2]. (b) Speed Profile of cells (n= 92) with terminal speed jump. (c) Speed Profile of cells (n= 60) with terminal speed jump. Red line shows the average instantaneous speeds whereas the gray shaded area shows the standard deviation.

## Discussion

CellMAPtracer is an open and easy-to-use software for tracking and extracting trajectory data of fluorescently-labelled cells through a user-friendly GUI. CellMAPtracer was designed with the aim to provide users with highly efficient tracking of migratory proliferating cells over multiple days through supervising, inspecting and correcting the tracking data in an enjoyable manner. As a proof of concept, breast cancer cells were scanned for 3 days. Over 100 cells were randomly tracked. To better evaluate and understand the resulting tracks, CellMAPtracer offers options to visualize and extract the resulting trajectory data. Users can interactively visualize any tracked cell and its descendants and compare the values of their migration measures and trajectory data (Fig. 1). Such comparisons give a quick and precise characterization of the tracked cells. In particular, it allows to unambiguously estimate the doubling time of the studied cells. Literature shows a wide spectrum in the doubling time of the BT549 cells, from 25.5 hours (Heiser et al., 2012), 51 hours (Cowley et la., 2014) to 3.7 days (Sweeney et al., 1998). A classical way of computing the doubling time uses initial and final cell counts in cultures and assumes exponential growth. CellMAPtracer, instead, enables user to get a real-time estimation of the doubling time directly from the trajectory time of the dividing daughter cells (Fig 3b). CellMAPtracer can also shed light on the synchronization degree in terms of of the division and migration between daughter cells. Our results for BT549 cells showed that two daughter cells do not follow the same migratory pattern and only 10% of the daughter cells divided synchronously. The speed-directionality dynamics prior and through cell division is an enigma. The categorical output of CellMAPtracer enables user with basic programming skills to gain extra insights about the speed-directionality dynamics of dividing cells. Our results showed that BT549 cells on 2D culture have during the G2-M phase have significantly less average speed, directionality and average instantaneous distance during the G2-M phase and 2D culture (fig. 6). A previous study (Esmaeili Pourfarhangi et al., 2018) reported such difference prior and through cell division in another breast cancer cell line cultured in 3D with no difference in 2D. During tracking, CellMAPtracer can help users detect unusual phenomena. We noticed an unusual phenomenon in BT549 cells, we refer it as the terminal speed jump we found in BT549 cells. A terminal speed jump was observed in 60.5% of the dividing BT549 cells. We also observed the phenomenon of multi-daughter cell division (i.e. more than two daughter cells), which is known to occur in aneuploid cancer cells. The three or more daughter cells were unevenly sized (Tse et al., 2012). While multi-daughter divisions are known, the terminal speed jump would be interesting to investigate. We can only speculate what this phenomenon might be attributed to. As cells progress towards and through mitosis, they are known to become rounder. This rounding is due to inactivation of the small GTPase Rap1 and consequently weakening or disassembling focal adhesions (Lancaster et al., Dev. Cell, 2013). The loosening of focal adhesions might enable cells to increase the speed for a very short period of time. Correlation of focal adhesion disassembly with the terminal speed jump is an interesting area for future investigation.

## Conclusion

Tracking migratory proliferating cells in long-term cultures with nearly 100% accuracy is a big challenge. Using CellMAPtracer, it is now straightforward to trace and follow individual cells and their descendants accurately. The lineage tracing of all descendant cells and their ancestors allows a better computation of the doubling time and understanding the heterogeneity of daughter cells and the speed-directionality dynamics prior and through cell division.

