## Supplementary Materials for "CellMAPtracer: A user-friendly tracking tool for long-term migratory and proliferating cells"

#### Supplementary Table 1

##### Available tracking tools

| Tool Name | Availability | Platform | Source code | Tracking | Accuracy | Performance | Cell division | Inspectability | Correctability | Categorized outcome |
|---|---|---|---|---|---|---|---|---|---|---|
| Braincells[1] | Free | Win | No | Auto | * | * | No | No | No | No |
| CellMAPtracer[2] | Free | Win/Lin/Mac | Yes | Semi | High | Moderate | Yes | Yes | Yes | Yes |
| CellProfiler[3] | Free | Win/Lin/Mac | Yes | Auto | Moderate | Fast | No | No | No | No |
| CellTrack[4] | Free | Win/Lin/Mac | No | Auto | * | * | No | Limited | Limited | No |
| CellTracker[5] | Free | Win | No | Semi | High | Moderate | No | Limited | Limited | No |
| ClusterTrack[6] | Free | Matlab | Yes | Auto | Moderate | Moderate | No | No | No | No |
| DcellIQ[7] | Free | Matlab | Yes | Auto | * | * | Yes | No | No | No |
| DIAS[8] | Paid | Win/Mac | No | Auto | * | * | No | No | No | No |
| DYNAMIK[9] | Free | Matlab | Yes | Auto | Moderate | Fast | Limited | No | No | No |
| FastTracks[10] | Free | Win | Yes | Auto | Moderate | Fast | No | No | No | No |
| ICY[11] | Free | Java | Yes | Auto | Moderate | Fast | No | No | No | No |
| ImarisTrack[12] | Paid | Win/Mac | No | Auto | * | * | Limited | Yes | Yes | No |
| LevelSetTracker[13] | Free | Matlab | Yes | Auto | * | * | No | Limited | Limited | No |
| LineageTracker[14] | Free | ImageJ | No | Auto | * | * | Yes | No | No | No |
| ManualTracking[15] | Free | ImageJ | Yes | Manual | Moderate | Very slow | Limited | Limited | Limited | No |
| MetaMorph[16] | Paid | Win | No | Auto | * | * | No | No | No | No |
| MTrackJ[17] | Free | ImageJ | Yes | Manual | Moderate | Very slow | No | No | No | No |
| NucliTrack[18] | Free | Win/Lin/Mac | Yes | Auto | High | Slow | Yes | Yes | Yes | No |
| ParticleTracker[19] | Free | ImageJ | Yes | Auto | Moderate | Fast | No | No | No | No |
| QuimP[20] | Free | ImageJ | No | Auto | * | * | No | No | No | No |
| TLA[21] | Free | Matlab | Yes | Auto | Moderate | Fast | No | No | No | No |
| tTt[22] | Free | Win | Yes | Semi | High | Moderate | Yes | Yes | Yes | No |
| Volocity[23] | Paid | Win/Mac | No | Auto | Moderate | Fast | No | No | No | No |

* Non tested tools

1. Hand, A. J., Sun, T., Barber, D. C., Hose, D. R., & MacNeil, S. (2009). Automated tracking of migrating cells in phase-contrast video microscopy sequences using image registration. Journal of microscopy, 234(1), 62-79.

2. Kamil Antos, & Salim Ghannoum. (2020, June 12). CellTracer 1.0 (Version v1.0). Zenodo. http://doi.org/10.5281/zenodo.3878088

3. Carpenter, A. E., Jones, T. R., Lamprecht, M. R., Clarke, C., Kang, I. H., Friman, O., Guertin, D. A., Chang, J. H., Lindquist, R. A., Moffat, J., Golland, P., and Sabatini, D. M. (2006). CellProfiler: Image analysis software for identifying and quantifying cell phenotypes. Genome Biol. 7, R100.

9. Mosig, A., Ja¨ger, S., Wang, C., Nath, S., Ersoy, I., Palaniappan, K. P., and Chen, S. S. (2009). Tracking cells in life cell imaging videos using topological alignments. Algorithms Mol. Biol. 4, 10.

10. DuChez B. J. (2017). Automated Tracking of Cell Migration with Rapid Data Analysis. Current protocols in cell biology, 76, 12.12.1–12.12.16. https://doi.org/10.1002/cpcb.28

11. de Chaumont, F., Dallongeville, S., and Olivo-Marin, J. C. (2011). ICY: A new opensource community image processing software. Proceedings of the IEEE International Symposium on Biomedical Imaging, pp. 234–237.

12. OMICS_06586, Bitplane, Switzerland

13. Dzyubachyk, O., Essers, J., Baldeyron, C., van Cappellen, W. A., Inagaki, A., Niessen, W. J., and Meijering, E. (2010b). Automated analysis of time-lapse fluorescence microscopy images: From live cell images to intracellular foci. Bioinformatics 26, 2424–2430.

14. Downey, M., Vance, K. W., & Bretschneider, T. (2011, March). Lineagetracker: A statistical scoring method for tracking cell lineages in large cell populations with low temporal resolution. In 2011 IEEE International Symposium on Biomedical Imaging: From Nano to Macro (pp. 1913-1916). IEEE.

15. Fabrice Cordelie´res (Institute Curie, Orsay, France)

16. Molecular Devices, USA

17. Meijering, E., Dzyubachyk, O., & Smal, I. (2012). Methods for cell and particle tracking. In Methods in enzymology (Vol. 504, pp. 183-200). Academic Press.

18. Cooper, S., Barr, A. R., Glen, R., & Bakal, C. (2017). NucliTrack: an integrated nuclei tracking application. Bioinformatics (Oxford, England), 33(20), 3320–3322. https://doi.org/10.1093/bioinformatics/btx404

19. Sbalzarini, I. F., and Koumoutsakos, P. (2005). Feature point tracking and trajectory analysis for video imaging in cell biology. J. Struct. Biol. 151, 182–195.

20. Bosgraaf, L., van Haastert, P. J. M., and Bretschneider, T. (2009). Analysis of cell movement by simultaneous quantification of local membrane displacement and fluorescent intensities using Quimp2. Cell Motil. Cytoskeleton 66, 156–165.

21. Huth, J., Buchholz, M., Kraus, J. M., Schmucker, M., Von Wichert, G., Krndija, D., ... & Kestler, H. A. (2010). Significantly improved precision of cell migration analysis in time-lapse video microscopy through use of a fully automated tracking system. *BMC cell biology*, *11*(1), 24.

22. Hilsenbeck, O., Schwarzfischer, M., Skylaki, S., Schauberger, B., Hoppe, P. S., Loeffler, D., ... & Strasser, M. (2016). Software tools for single-cell tracking and quantification of cellular and molecular properties. Nature biotechnology, 34(7), 703-706.

23. PerkinElmer®, USA

### Supplementary file S1

##### Cell Culture

BT549 cells were cultured at 37°C and 5% CO2 in RPMI supplemented with 10% FBS (Fetal Bovine Serum) and 100 U/ml penicillin/streptomycin. To generate GFP-BT549 stably-labeled cells, BT549 cells were transduced (according to the Essen BioScience protocol) with NucLight lentivirus. The green protein-based fluorophores of NucLight are located within the nuclear envelope of the cell. During cell division new NucLight protein is synthesized and transmitted to the new daughter cells. The transduced cells were sorted via fluorescence-activated cell sorting (FACS) using a BD FACSAriaTM cell sorter. For optimal tracking efficiency a mixed population of BT549 and GFP-BT549 cells were used for the random migration assay. These cells were co-cultured at a ratio of 1:3 respectively.

##### 2D random migration and live cell imaging

15000 cells from the mixed population were cultured per well in 96 well image-lock plate (EssenBio, 4739, Lot#17040501) for 24 hours at 37°C and 5% $CO_2$. And then cells were scanned at ten-minute intervals over three days in Essen BioScience's IncuCyte S3 using both the phase and green channels. Cellular viability was assessed throughout the course of the scanning by comparing the phase cellular morphology between BT549 and GFP-BT549 cells .

### Supplementary fig.1

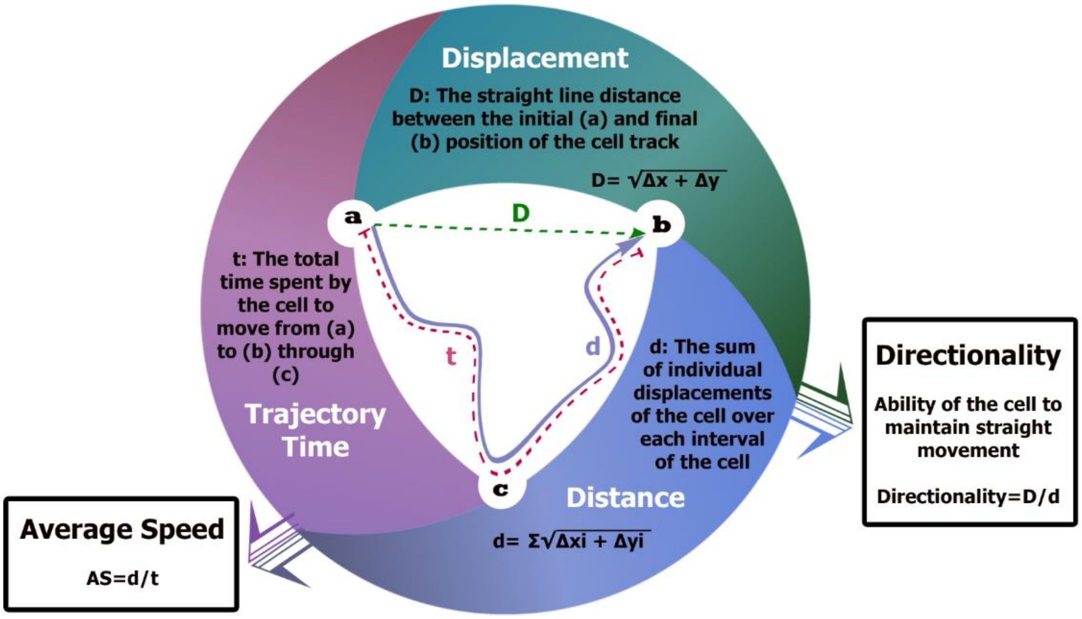


*Supplementary Fig. 1: A schematic diagram elucidates how the migration measures are calculated in addition to description of the quantification acquired.*

### Supplementary fig.2

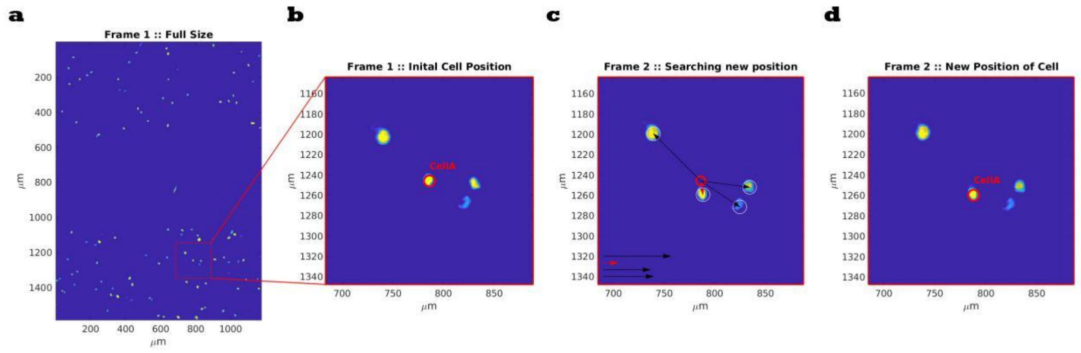

*Supplementary Fig. 2: A schematic diagram explaining the tracking paradigm. (a) A full size view of the first frame of a multi-frame tiff file. (b) A selected field of the first frame of the multi-frame TIFF file showing nuclei of 4 cells, the red ring refers to the location of the target cell nucleus. (c) Distances between the last position of the target cell nucleus and the position of each nucleus in the second frame, red arrow refers to the shortest distance. (d) new position (red ring) of the tracked cell.*

### Supplementary fig.3

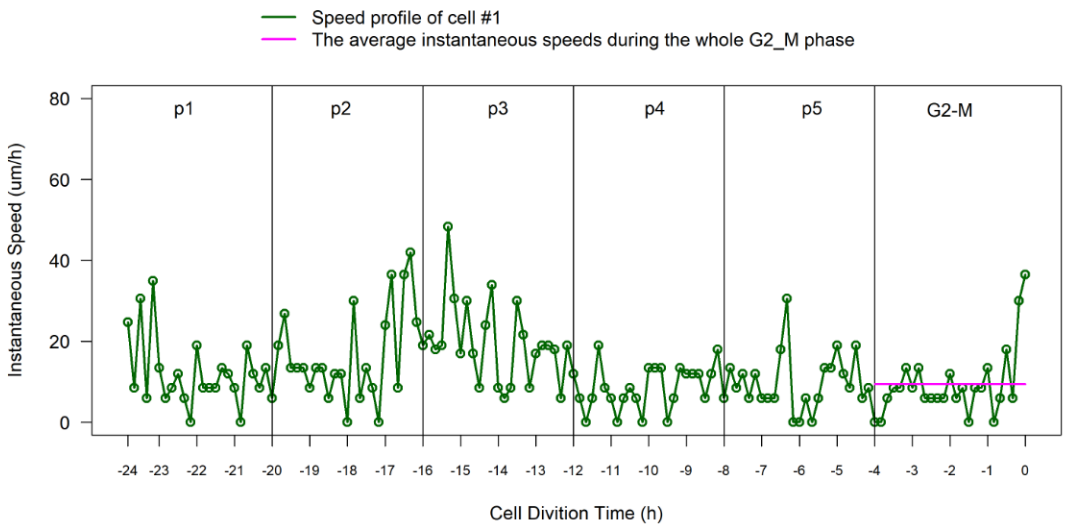


*Supplementary Fig. 3: Speed profile*
